# Natural history of a mouse model of X-linked myotubular myopathy

**DOI:** 10.1101/2021.10.11.463940

**Authors:** Ege Sarikaya, Jonathan Volpatti, Nesrin Sabha, Nika Maani, Hernan D. Gonorazky, Alper Celik, Paula Onofre-Oliveira, James J. Dowling

## Abstract

X-linked myotubular myopathy is a severe monogenetic disorder of the skeletal muscle caused by loss of expression/function mutations in the *MTM1* (myotubularin) gene. There is a growing understanding of the pathologic and molecular abnormalities associated with loss of MTM1, and emerging therapeutic strategies that are in the process of translation to patients. Much of these data have been uncovered through experimentation in pre-clinical animal models of the disease. The most widely used model is an Mtm1 gene knockout mouse line; this line faithfully recapitulates the salient genetic and pathologic features of the disease. Despite the advances in aspects of XLMTM, there remain many unknowns related to disease pathomechanisms and to understanding of MTM1’s function in normal muscle development, and a continued need for therapy identification and development. To address these barriers, and to lay the groundwork for future study, we performed a natural history study of the Mtm1 knockout mouse model of XLMTM. We show that certain molecular and pathologic changes precede overt phenotypic changes, while others, including abnormalities in triad structure, occur more coincident with muscle weakness in the mouse. In total, we provide a comprehensive longitudinal assessment of molecular and structural features of the murine XLMTM disease process.

## Introduction

X-linked myotubular myopathy (or XLMTM) is a monogenetic muscle disease with onset in infancy characterized by neonatal hypotonia and severe weakness(Lawlor and Dowling, 2021). Between 25-50% of affected patients die in the first year of life, and those that survive have a high degree of technology dependence (80% requiring wheelchair, ventilator, and feeding tube support) and shortened life span(Amburgey et al., 2017; Annoussamy et al., 2019). Historically, the condition was diagnosed by characteristic features on muscle biopsy including centrally located myonuclei, myofiber hypotrophy, and accumulation of mitochondria and other perinuclear organelles. Currently, diagnosis is established with positive genetic testing(Gonorazky et al., 2018). Of note, female carriers are often asymptomatic, though there is a growing recognition of a range of manifesting phenotypes(Cocanougher et al., 2019; Reumers et al., 2021).

XLMTM has a single genetic cause, i.e. mutations in myotubularin (or *MTM1*)(Laporte et al., 1996). Mutations in *MTM1* are all thought to result in loss of expression and/or loss of function(McEntagart et al., 2002). MTM1 encodes the myotubularin (MTM1) protein, which is a lipid phosphatase that acts on phosphoinositides (PIPs)(Hnia et al., 2012). Specifically, MTM1 dephosphorylates (and thus deactivates) PI3P and PI3,5P2(Volpatti et al., 2019). Through this action, it also generates PI5P. MTM1 is a resident endosomal protein, and its main role in cells is to regulate vesicular sorting through the endosomal compartment via a PIP conversion mechanism(Ketel et al., 2016).

Several pathologic and molecular abnormalities have been identified in muscle from MTM1 deficient patients and animal models. In terms of pathologic alterations, in addition to the structural changes typical observed on muscle biopsy, important abnormalities in the appearance and function of the excitation-contraction (EC) coupling apparatus have been described(Al-Qusairi et al., 2009; Dowling et al., 2009). EC coupling is the process via which signals initiated at the neuromuscular junction are converted to muscle contraction(Dowling et al., 2021). The key muscle substructure that mediates this process is the triad, which represents the apposition of the T-tubule (a membrane invagination that contains the dihydropyridine receptor) and the terminal sarcoplasmic reticulum (which contains the ryanodine receptor, which is a calcium channel and main effector of EC coupling). In XLMTM, the structure of the triad is abnormal, and EC coupling is abnormal; these changes are thought to be responsible for the weakness observed in patients(Jungbluth et al., 2018).

Several molecular abnormalities have been described. These include changes in the AKT pathway, aggregation of desmin intermediate filaments(Hnia et al., 2011), increased levels of the large GTPase DNM2(Cowling et al., 2014), altered protein ubiquitination(Gavriilidis et al., 2018), and changes in autophagic flux(Al-Qusairi et al., 2013; Bachmann et al., 2017; Fetalvero et al., 2013). Some of these abnormalities are likely the result of direct interactions with MTM1, which has been shown to interact with desmin and with UBQLN2. Of these changes, perhaps the most importance is increases in DNM2 protein levels, as this has been shown to directly lead to many of the pathologic changes characteristic of XLMTM(Cowling et al., 2011; Liu et al., 2011). In this regard, therapies aimed at lowering DNM2 (including antisense oligonucleotide (ASO) mediated knockdown and chemical reduction with tamoxifen) have shown great efficacy in pre-clinical models and currently in clinical trial in patients(Maani et al., 2018; Tasfaout et al., 2017). In addition, gene replacement therapy can rescue the phenotype of all pre-clinical models and is also in clinical trial(Childers et al., 2014).

Much of what is known about XLMTM has been uncovered through experimentation in a mouse model of the disease. This model, first generated by Buj-Bello and Laporte(Buj-Bello et al., 2002), has a targeted deletion of exon 4 of murine *Mtm1*, the consequence of which is frameshift, reduced RNA expression, and absent protein expression (i.e. a gene knockout or KO model). Other animal models have also been established, including zebrafish and canine models(Beggs et al., 2010; Dowling et al., 2009; Sabha et al., 2016), that harbor mutations in the MTM1 gene. All vertebrate models tested have faithfully recapitulated the major abnormalities seen in patient muscle, including central nuclei, myofiber hypotrophy, and triad disorganization, and are associated with abnormal motor function and early lethality.

Much remains to be understood about the underlying pathomechanisms of XLMTM. In particular, the link between MTM1’s role as a phosphoinositide phosphatase and regulator of endosomal dynamics and the pathologic and molecular changes in XLMTM muscle are not clear. Furthermore, it is not known what the first/inciting events are in the disease process. In addition, as more therapies are identified for the disease, it is critical to understand how these treatments may synergize, and how they compare in terms of effectiveness for ameliorating key aspects of the disease. To begin to address these unknowns, we have undertaken to define the natural history of the XLMTM mouse model. By evaluating pathologic and molecular events at different time points, we identify that mitochondrial changes, myofiber hypotrophy and increased DNM2 are among the earliest changes observed. Surprisingly, triad changes are found later in the disease, process, after a period of normal triad appearance. In total, our work provides critical groundwork for future studies related to therapy testing and establishment of disease pathomechanisms.

## Results

### Mtm1 knockout (KO) mice have four stages of disease

We have previously established a colony of *Mtm1* KO mice on the C57BL6J background(Maani et al., 2018; Sabha et al., 2016). We have observed 4 phases of disease in the mice (Figure 1A). The first is the pre-symptomatic phase, which occurs until the age of weaning (21 days postnatal). For analysis purposes, we have chosen age 14 days to represent this time period. The second is symptom onset, which occurs at approximately 21 days postnatal. The first observed phenotypic abnormality is reduced body weight (Figure 1B). The third is onset of motor deficits, which in our hands occurs at approximately 28 days of age. The last is the endpoint phase, where KO mice have severe hindlimb paralysis and generalized wasting and typically require termination. This occurs at 35 days and beyond (median survival 38 days).

**Figure 1:**
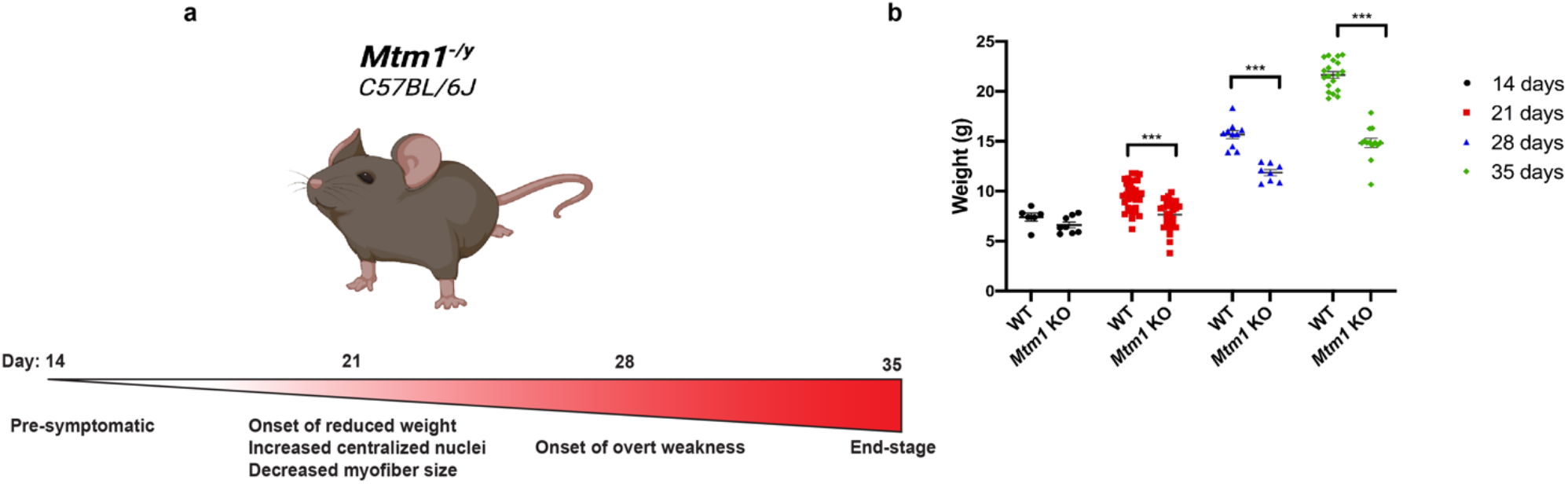
natural history of the *Mtm1* knockout mouse model. (A) Schematic depicting the 4 phases of the disease process in *Mtm1* knockout (*Mtm1*^*-/y*^) mice. (B) Scatter plot comparing the weight (g) of WT and Mtm1 KO mice at 14 (WT n= 6, KO n=8), 21 (WT n=30, KO n=26), 28 (WT n=10, KO n=8) and 35 (WT n=19, KO n=13) days. Average weight of WT mice is 7.40g, 9.18g, 15.65g and 21.64g, at 14, 21, 28 and 35 days respectively and average weight of KO mice is 6.61g, 7.59g, 11.86g and 14.83g, at 14, 21, 28 and 35 days respectively. (***p<0.001).

### Mitochondrial and sarcolemmal disorganization represent the first structural changes

We initiated our study by analyzing muscle histopathology at our 4 defined stages. We specifically evaluated changes that have previously been described in *Mtm1* KO mice, which include % central nuclei and myofiber size (both quantified from hematoxylin and eosin staining), mitochondrial distribution (defined by succinate dehydrogenase staining), sarcolemmal membrane organization (illuminated by dysferlin immunostaining), and triad structure (defined by electron microscopy) (Supplemental Tables). We performed these analyses on cryosections of skeletal muscle at 14, 21, 28, and 36 days postnatal on KO vs WT littermates, using a minimum n = 4 per genotype with blinded examination.

Interestingly, despite the fact that mice are overtly normal at 14 days of age, muscle structure is clearly altered at this stage (Figure 2A). The most striking finding is redistribution of mitochondria into “ring” fibres, something that is seen on occasion in WT 14 day olds but that is significantly increased in KO mice (average of 1.0 per field for WT vs 6.7 for KO). There are also small but significant changes in % central nuclei and dysferlin immuno-localization. While the distribution of fiber size is altered, mean myofiber diameter is not significantly changed (data not shown). In addition, triads appear normal (data not shown). It is notable that none of the changes occur in more than a minority of fibers.

**Figure 2:**
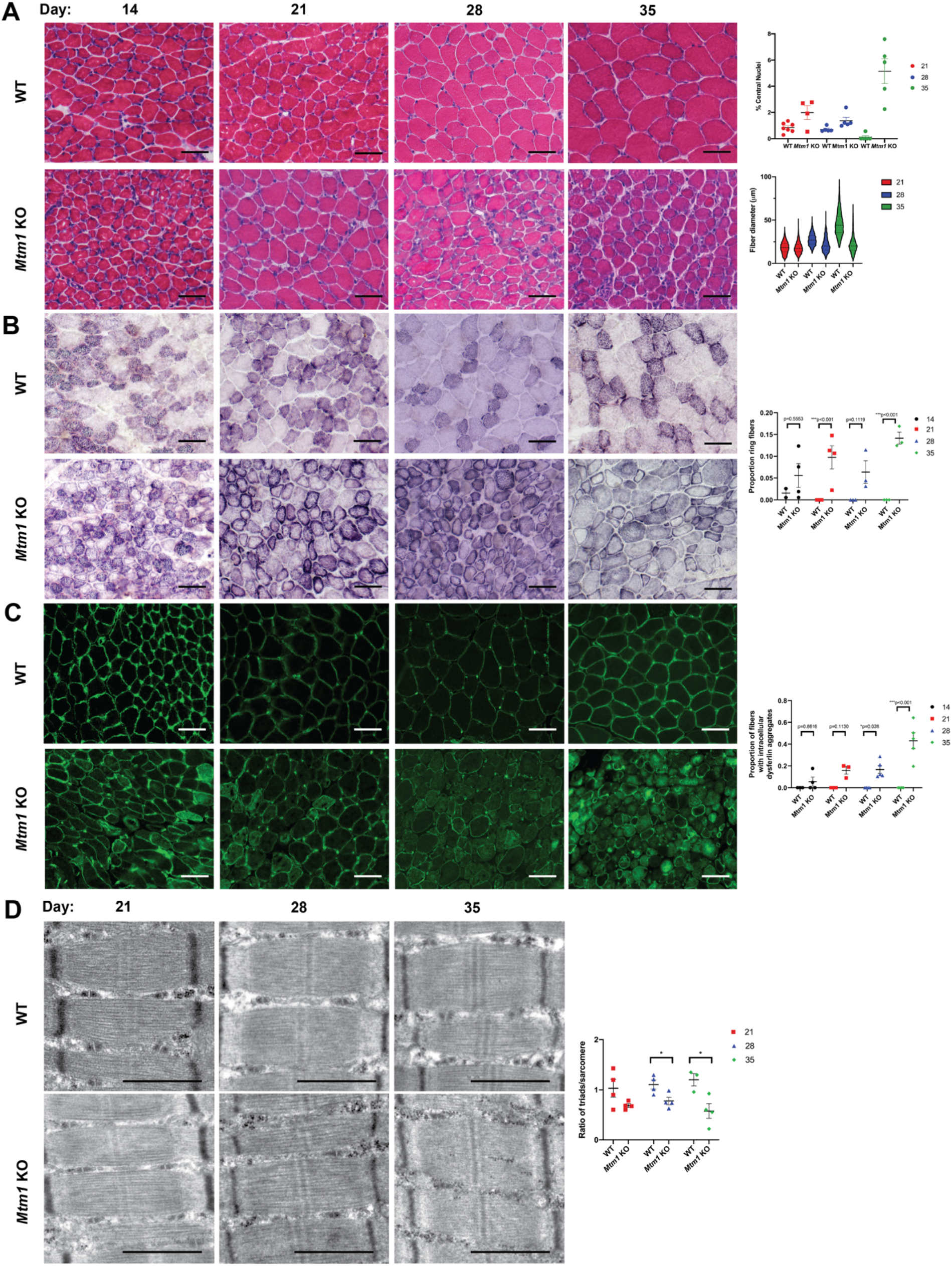
Progressive histopathologic changes in *Mtm1* knockout mice. Histopathology as defined by staining with hematoxylin/eosin (A), succinate dehydrogenase (B), and anti-dysferlin immunofluorescence (C) on cryosections and electron microscopy on ultrathin sections (D). Subtle changes in SDH staining and dysferlin localization are noted at 14 and 21 days. By 21 days, there are also increases in % central nuclei. Myofiber hypotrophy is present at 21 days in a small number of fibers, with fiber size difference substantially increasing in magnitude at 28 and 35 days. Significant reduction in triad number and abnormal triad appearance is observed late in the disease process (i.e. at 28 and 35 days), coincident with overt muscle weakness. Scale bars (A-C) = 50 um and (D) = 1 um. **** p = 0.0001

At 21 days, these abnormalities are all more pronounced (Figure 2A-C), but triad structure remains unchanged (Figure 2D). In fact, there is no quantitative difference in the number or appearance of triads at this age It is not until 28 days of age that triads become statistically different in KOs vs WTs (Figure 2D). By 36 days of age, there are very few muscle fibers that appear normal (Figure 2A-D).

### Muscles are differentially effected in Mtm1 KO mice

In XLMTM, there is variability in the extent of pathology seen in different muscle groups. This is best appreciated in patients by using muscle MRI(Carlier and Quijano-Roy, 2019). We examined this in our *Mtm1* KO mice, using both MRI and matched histopathological analysis. We studied mice at 35 days of age, which was optimal age for MRI scanning in terms of mouse size. Using T2 weighted MRI images, we defined muscle volume of the muscle in the distal hindlimb. We found that plantaris and tibialis anterior (AT) were the most affected in terms of reduced size, while the soleus and tibialis posterior were the least affected (Figure 3B). To relate these findings to standard muscle pathology, we measured % central nuclei and myofiber size from H/E stained cryosections of muscle taken from the same mice that we imaged (Figure 4). We saw good correlation between MRI findings and % central nuclei, with plantaris and TA having the highest %. Plantaris also displayed the largest magnitude change in myofiber size, with TA also decreased substantially. Conversely, extensor digitorum longus (EDL), which was relatively mildly affected per MRI, had significant changes in myofiber size and % central nuclei.

**Figure 3.**
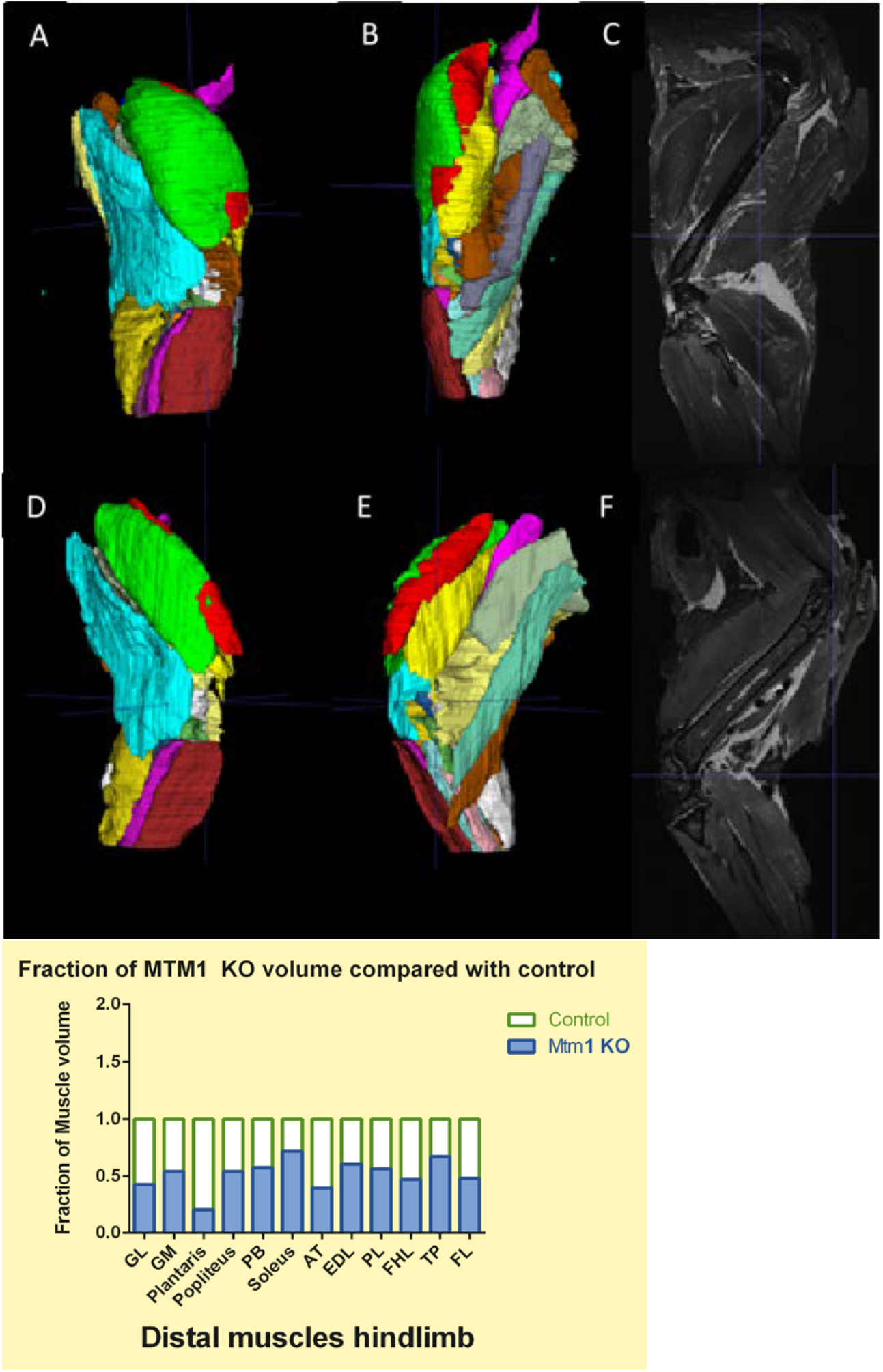
MRI of skeletal muscle from wild type and Mtm1 KO male mice. Muscle MRI was done on post-mortem mice at age 35 days. Limbs were fixed and prepared by a protocol described elsewhere. T1 and T2 weighted images were obtained. Depicted (top panel) are 3D reconstructions of right hind limbs of wild type (A,B,C) and MTM mice (D,E,F). The segmentation was done manually using the software ITK-scan. (Bottom panel) Volumes of all muscle were decreased in Mtm1 KO mice, but there was important variability between different muscle groups. Volumes are expressed as a fraction of wild type (control), with wild type normalized to a value of 1.

**Figure 4.**
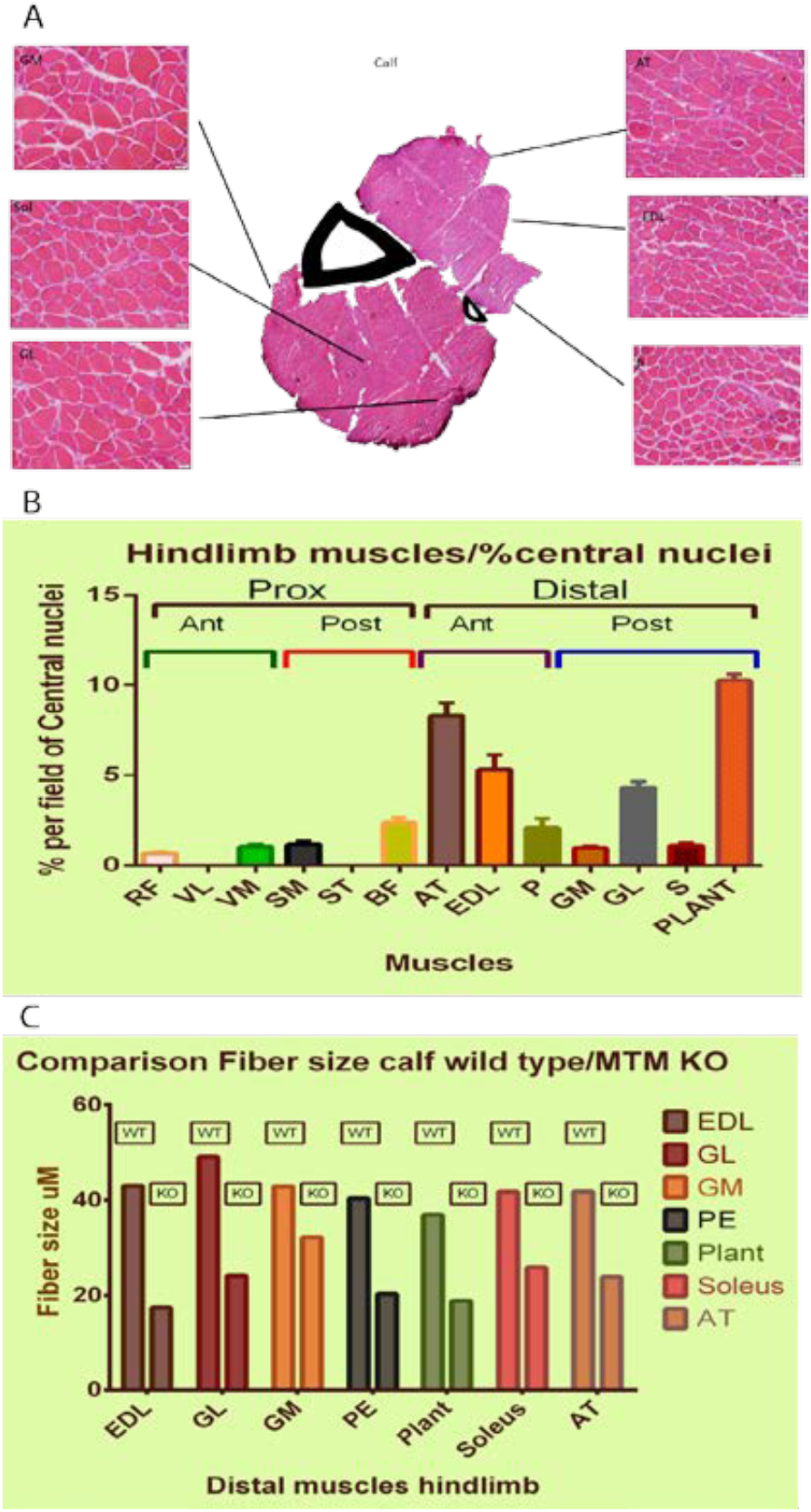
Comparison of histopathologic characteristics between muscle groups. (A) Histology of the distal portion of the hind limb with analysis of each of the muscle of the Mtm1 KO mice. Each portion was dissected and prepared for cryosectioning. Each muscle was analyzed for muscle fiber size and % of central nuclei. (B) % of central nuclei of the hindlimb muscles. Distal muscles were most impacted, with AT, EDL, and PLANT with the highest values. (C) The most affected muscles in terms of fiber size were Extensor digiti longs (59% reduction in size as compared to wild type), plantaris (50.7%), gastrocnemius lateralis (50.6%) tibialis anterior (42.83) (RF rectus femoralis, VL vastus lateralis, VM vastus medialis, SM semimembranosus, ST semitendinosus, BF bicep femoralis, AT tibialis anterior, EDL extensor digitalis longus, P plantaris, GM gastrocnemius medialis, GL gastrocnemius lateralis, S soleus, P plantaris)

### Increases in DNM2 protein levels and acetylated tubulin are among the first molecular changes observed in Mtm1 KO mice

We next examined by western blot several of the molecular changes that have been described in *Mtm1* KO mice (Figure 5). These include DNM2 levels, polyubiquitination, autophagy (as indicated by p62 levels), and acetylated tubulin. At 14 days postnatal, only DNM2 and tubulin were significantly changed as compared to WT. The magnitude of change, however, was relatively small. At 21 days, all markers examined were abnormal, with changes increasing in their differences vs WT at each age.

**Figure 5:**
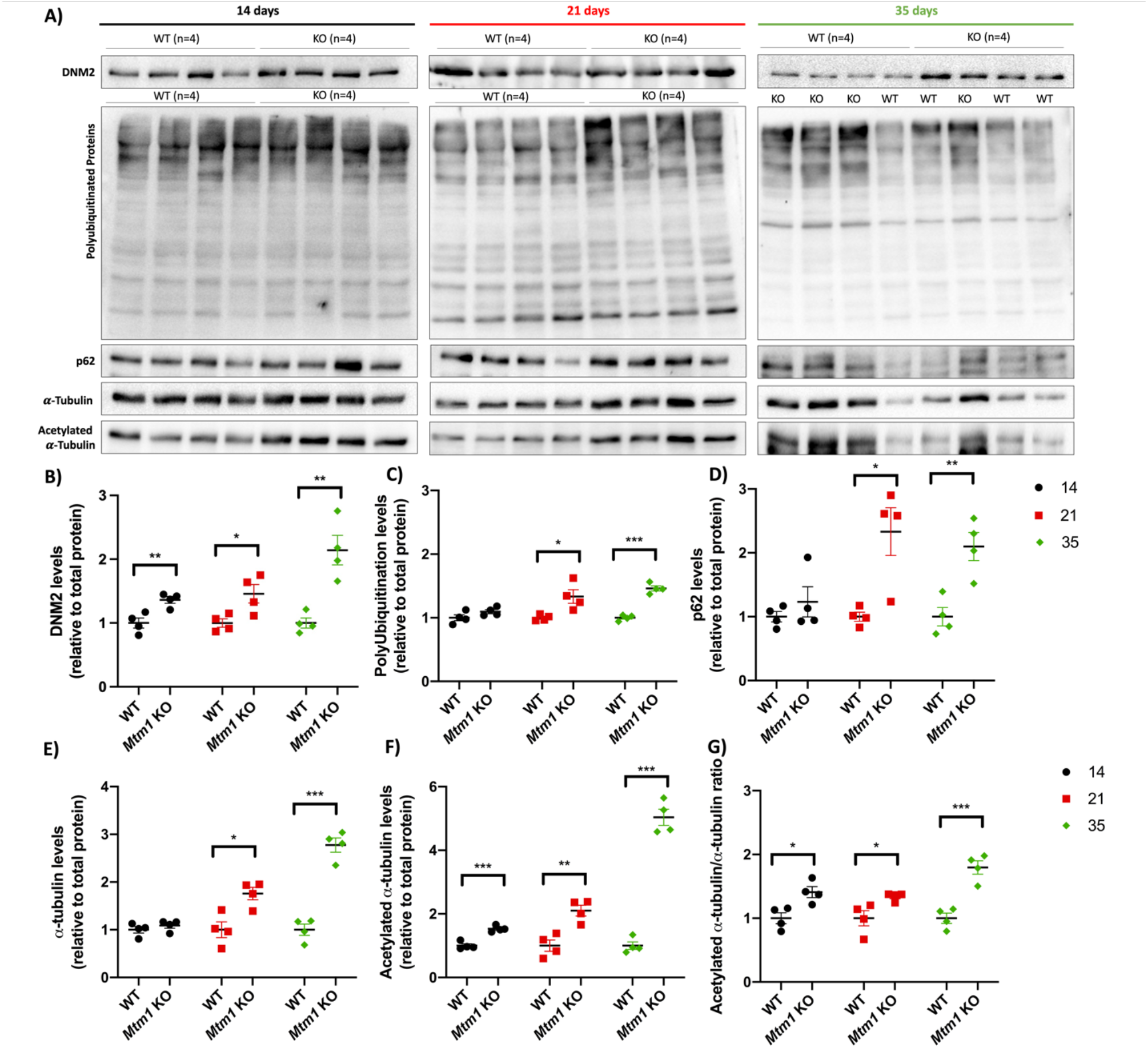
Molecular changes in *Mtm1* knockout mice. A) Representative western blot images of DNM2, polyUbiquitination, p62, *a*-tubulin and acetylated-*a*-tubulin at 14, 21 and 35 days. B-G) Quantification of protein levels relative to total protein staining and normalized to average WT expression (see also Supplemental Figure 1). B) DNM2 levels are elevated at 14 (**p=0.0085), 21 (*p=0.0287) and 35 (**p=0.0034) days. C) Polyubiquitinated protein levels are elevated at 21 (*p=0.0228) and 35 (***p<0.001) days. D) p62 levels are elevated at 21 (*p=0.0126) and 35 (**p=0.0058) days. E) *a*-tubulin levels are elevated at 21 (*p=0.0115) and 35 (***p<0.001) days. F) Acetylated-*a*-tubulin levels are elevated at 14 (***p<0.001), 21(**p=0.0041) and 35 (***p<0.001) days. G) Acetylated-*a*-tubulin/*a*-tubulin ratio is elevated at 14 (*p=0.0157), 21 (*p=0.0339) and 35 (***p<0.001) days. All quantifications included n=4/genotype, and 3 technical replicates.

### Comparative RNA sequencing identifies emerging pathway abnormalities

To identify molecular pathways altered with *Mtm1* KO, we performed bulk RNA sequencing on anterior tibialis muscle at 14, 21, and 35 days (Figure 6). Very few transcriptional changes were noted at 14 days of age, with only 16 differentially expressed genes (DEGs) at this time point (Figure 6A). By 21 days, however, there are widespread alterations in gene expression, with more than 800 DEGs. By 35 days of age, >2500 DEGs are identified, with numerous cellular pathways disturbed. Principal component analysis shows that the difference between genotypes increases with age (Figure 6D).

**Figure 6.**
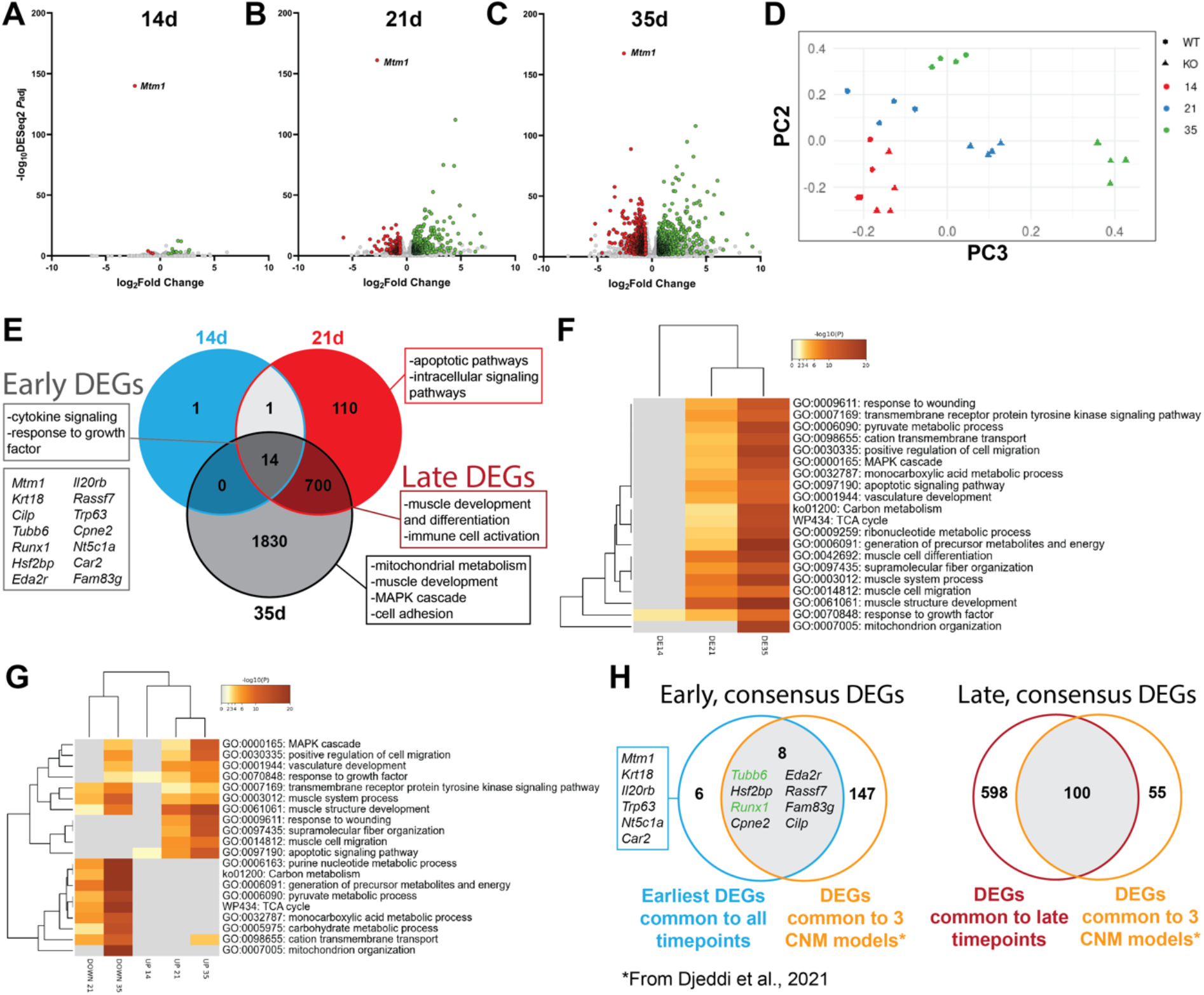
Longitudinal transcription changes in *Mtm1* knockout mice. A-C) Volcano plots highlighting differentially expressed genes between WT and *Mtm1* KOs from tibialis anterior (TA) muscle at A) 14 days, B) 21 days, and C) 35 days (Adjusted p-value<0.01, log2FC>0.585). D) Principal component analysis plot of all timepoints indicating gene expression is similar between WT and KO mice at the earliest stage tested but diverges starkly from 21 days on. E) Venn diagram depicting common and unique differentially expressed genes at each timepoint. Boxes highlight major GO enrichment terms with the associated DEG list. F-G) Heat map visualization of GO terms enriched at each timepoint considering F) all DEGs in each timepoint and G) by up- or down-regulated status. H) Venn diagram highlighting shared DEGs between our dataset and those identified as being in common in three centronuclear myopathy mouse models (including the same *Mtm1* KO mice on a different background) by Djeddi and colleagues.

We considered which pathways were affected at each stage by performing gene ontology (GO) term enrichment (Figure 6E-G). The few DEGs at 14 days were associated primarily with apoptotic pathways and growth factor signaling. By 21 days, many dysregulated pathways emerge, including down-regulation of mitochondrial metabolism genes and elevated expression of inflammatory pathways (wound response and apoptotic signaling), and dysregulation of intracellular signaling pathways. Finally, at the end-stage, we find MAPK pathway genes are strongly dysregulated and that there is further dysregulation of mitochondrial pathways.

Lastly, we examined for specific DEGs found in common with a recent transcriptomic study comparing three centronuclear myopathy models, including the *Mtm1* KO mice (Figure 6H)(Djeddi et al., 2021). This comparison revealed eight consensus DEGs changed early in the disease process. For example, *Runx1* is upregulated the earliest time point investigated, prior to observable muscle weakness. Additionally, *Tubb6* was upregulated in all data sets, and also at all time points in our study.

### Subcellular proteomics identifies changes in the proteosomal network and shows that the majority of changes occur in the organelle and membrane fraction

To corroborate our gene expression findings, and to better understand the subcellular relationship of the molecular changes, we performed proteomics on subcellular fractions of 21 days old skeletal muscle. We first confirmed the success of our fractionation methodology (Figure 7A) by examining of expression of proteins known to localize to different compartments (Figure 7B). In general, we saw good segregation between the 5 fractions generated. Notably, the nuclear fraction exhibited the most contamination by sarcomeric/myofibrillar components. We then explored differences between WT and KO mice, and found the most significant differences in the membrane/organelle fraction, where > 200 proteins were differentially expressed (DEPs). Interrogation of pathways associated with these DEPs identified fiber organization, vesicle transport, and focal adhesions as among the most significantly changed.

**Figure 7.**
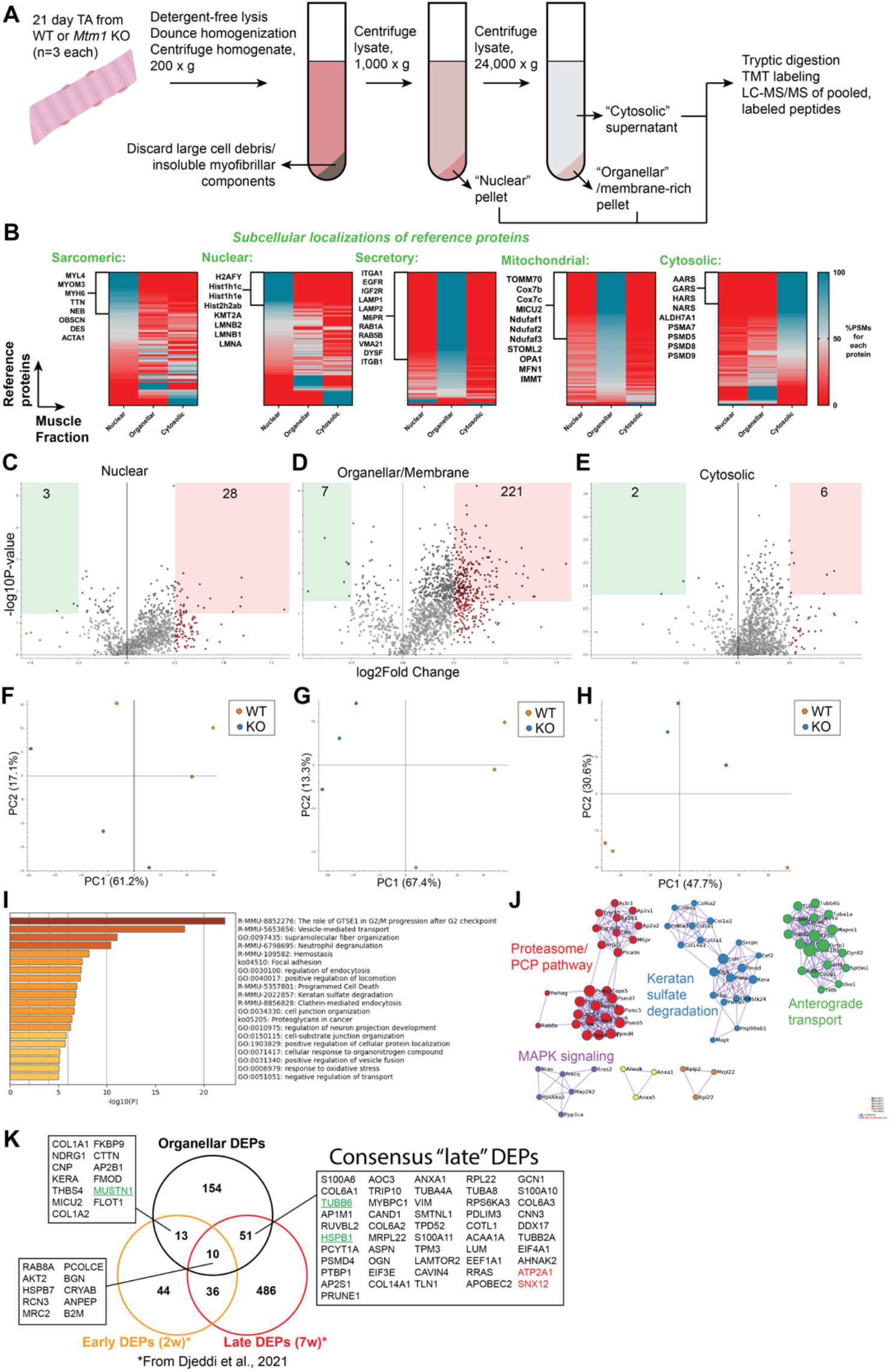
Subcellular proteomics in *Mtm1* KO mice. A) Overview of methods used to perform proteomics on three subcellular fractions from 21 day mouse tibialis anterior (TA) muscle. See Methods section for details. B) Heat maps representing the enrichment of proteins in each of the muscle fractions with reference to consensus subcellular location data. Peptide spectrum matches (PSMs) provide an estimate of protein abundance and were used to show what percentage of PSMs for a given protein were identified in each fraction. Data are sorted based on which fraction had the greatest number of 100% PSMs for proteins in the reference set. C-E) Volcano plots highlighting differentially expressed proteins (DEPs) in the C) nuclear, D) organellar/membrane, and E) cytosolic fractions (p-value<0.05, log2FC>0.5). F-H) Principal component analysis plots from each fraction. I) GO term enrichment analysis of DEPs from the organellar/membrane fraction reveals dysregulation of focal adhesions and vesicle-mediated and endocytic pathways in *Mtm1* KOs. J) Protein-protein interaction network visualization of organellar fraction DEPs show aberrant levels of proteasomal proteins and MAPK signaling pathways in *Mtm1* KOs. K) Venn diagram highlighting DEPs found in common between the organellar fraction and those identified in a longitudinal study of *Mtm1* KO mice by Djeddi and colleagues. Proteins in green were also identified as either early or late consensus DEGs. All DEPs shown had the same “sign”, i.e. up-versus down-regulated, except for the two proteins indicated in red.

We performed network analyses from these data as well, and saw changes in the UPS and anterograde transport among others. Of note, the membrane/organellar fraction was enriched in mitochondrial proteins. Given that mitochondrial abnormalities were observed at the 21 day time point by both histology and by RNAseq, we focused on mitochondrial DEPs. We found levels of MICU2, ATP5F1C, PDP1, and MRPL22 were elevated while NDUFA3, CYCS, and MT-CYB were decreased, providing further support for dysregulation of mitochondrial function. Lastly, we highlight specific proteins that were also differentially expressed in a recent study using bulk proteomics in the *Mtm1* KO mice(Djeddi et al., 2021).

### MTM1 interacts with both MTMR10 and MTMR12 as well as several novel proteins

Lastly, we sought to identify relevant protein-protein interactions for MTM1 at different time points. To accomplish this, we used CRISPR/Cas9 gene editing to introduce a 6x-HIS tag in frame with the 3’ end of the MTM1 gene. We then performed immunoprecipitation followed by mass spectrometry using anti-His beads (Figure 8A). We started with muscle extracts from 35 day old mice (Figure 8B). We identified two known interactors, BIN and MTMR12, validating the ability of our methodology for finding MTM1 binding partners(Gupta et al., 2013; Royer et al., 2013). We also identified MTMR10 which, like MTMR12, is a phosphatase dead member of the myotubularin family. Interestingly, MTMR10 localizes both to the cytoplasm and the nucleus, suggesting the possibility that some MTM1 may be found outside of the endosome. This is further suggested by some of the other novel binding partners such as reticulon 2 (RTN2) and Nuclear Receptor Subfamily 4 Group A Member (NR4A3). We also identified IMMT, a member of the mitochondrial contact site and cristae organizing system (MICOS) complex, that has previously been identified as an MTM1 interactor in two independent studies in HEK293 cells. This finding suggests direct interplay between MTM1 and the mitochondria and goes along with the pathologic and gene expression changes seen in mitochondria with loss of *Mtm1*. We then performed a similar analysis on muscle extracts from 21 day old mice (Figure 8C). Surprisingly, we found essentially a non-overlapping group of proteins. Of note, there were still several mitochondrial proteins identified, further strengthening a direct link with mitochondria. There were also multiple associations with regulators of nucleotide metabolism.

**Figure 8.**
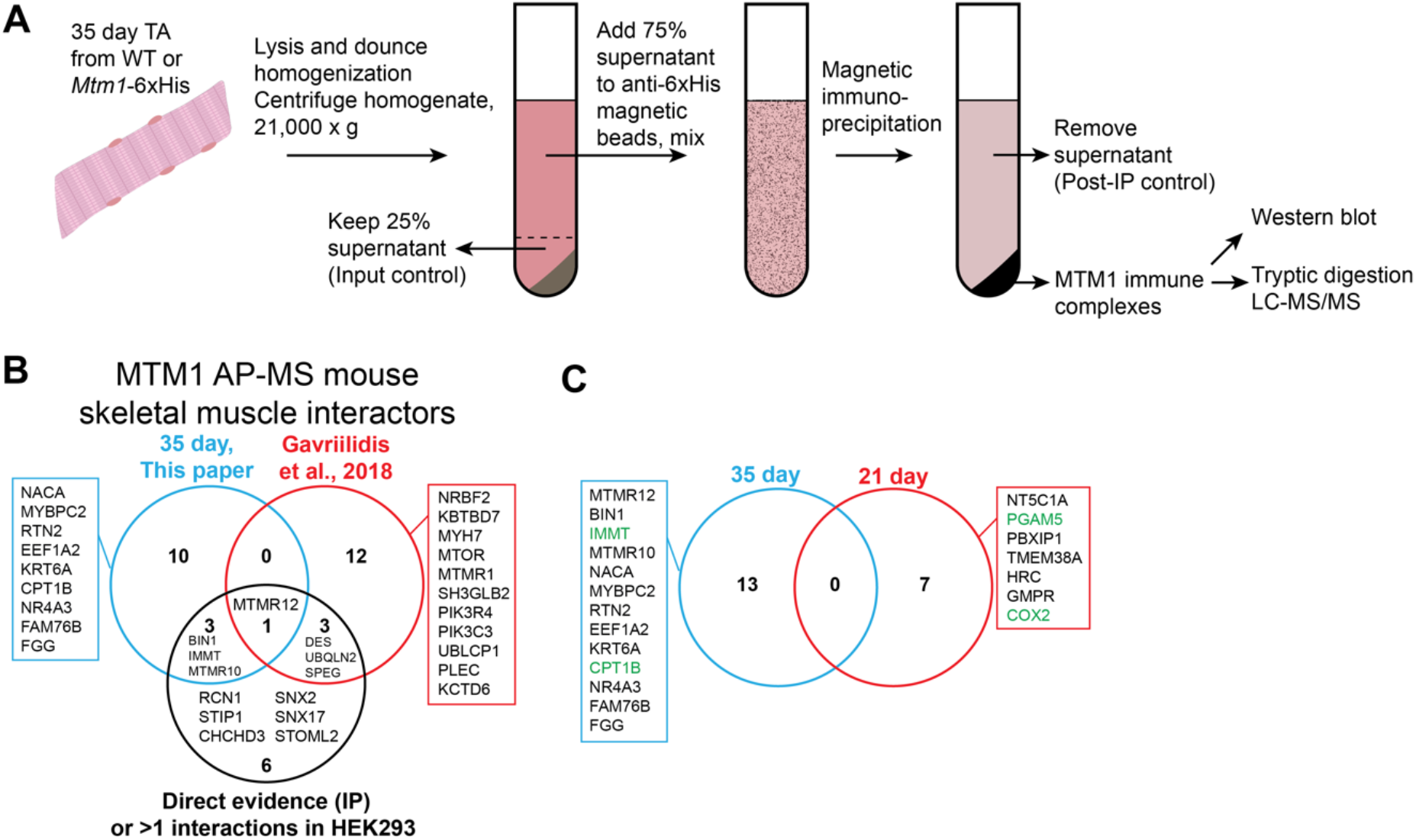
MTM1 protein interactions in skeletal muscle at 21 and 35 days. (A) Methodology for identifying putative MTM1 interactors using Mtm1-6xHis tagged mice and immuno-precipitation followed by mass spectrometry. (B) Interactors identified from skeletal muscle extracts at 35 days, with comparison to interactors identified in previous studies. (C) Interactors from 21 day muscle extracts. Note no overlap with the proteins identified at 35 days.

## Discussion

In this study, we present the natural history of a mouse model of X-linked myotubular myopathy (Figure 9). We identify molecular and structural changes in the muscle that precede the onset of overt phenotypic abnormalities, showing that the disease process occurs during muscle development and before the animal’s present outward signs of weakness and muscle dysfunction. These early changes do not include alteration of triad structure, showing that triad formation occurs normally and suggesting triad abnormalities are either a secondary consequence or the result of loss of MTM1 function in triad maintenance. These and other observations lay the groundwork for future experimentation aimed at undercovering the role of MTM1 in normal muscle development and elucidating the first steps in the XLMTM disease pathomechanism. Finally, our data provide a guide for harmonization and uniformity of future pre-clinical interventional studies.

**Figure 9:**
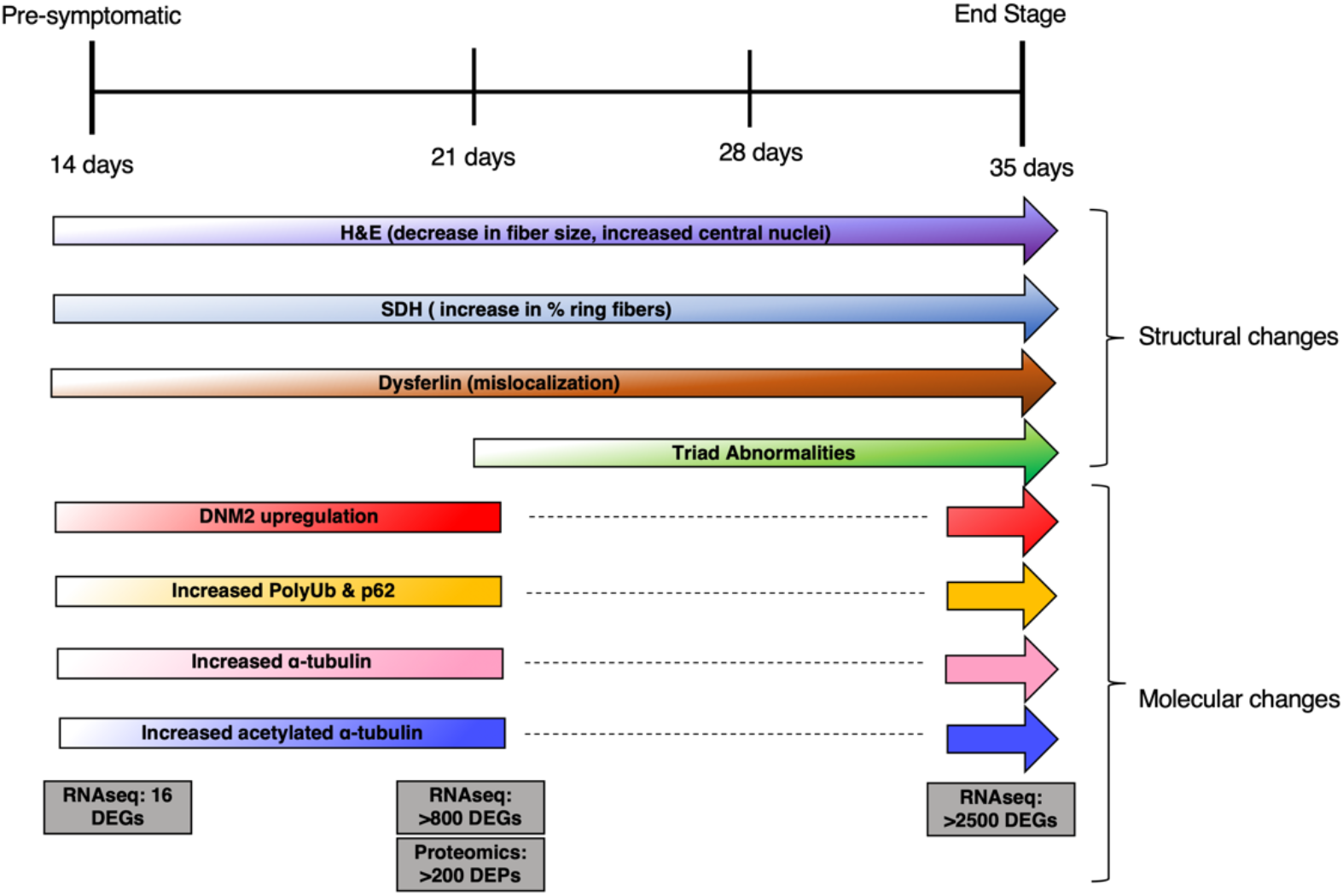
Summary of pathologic and molecular changes in the *Mtm1* knockout mouse model. Summary of the findings from this manuscript, including changes in histopathology and in molecular markers, mapped across the 4 stages of the murine disease process.

One of the most striking observations in our study is the early and pervasive change in mitochondrial distribution. This appears to be the first structural change in the muscle, and is consistent with the known interplay between MTM1 and desmin, as the desmin intermediate filament network is known to be a key regulator of mitochondrial organization. MTM1/mitochondrial interplay may additionally be mediated by interactions between MTM1 and mitochondrial components, as suggested by our IP-MS data, which identified several mitochondrial proteins. In total, there is an intriguing possibility that MTM1 may be a key mediator of mitochondrial tethering, a process important for mitochondrial distribution and function and controlled in part via interaction with the endosome. In terms of the disease process, mitochondrial disorganization may contribute to changes in myofiber size and thus to overall body weight. It may also influence other molecular pathways that are abnormal later in the disease process, such as autophagy and the ubiquitin-proteasome system.

In contrast, triad changes do not occur until later in the disease process. They do appear to coincide with overt muscle weakness, consistent with the hypothesis that they are a main driver of muscle weakness in XLMTM. The interplay between MTM1 and the triad remains poorly understood. Our findings do not necessarily shed new light on this, except to support the existing idea that MTM1 is not required for triad formation, but rather for triad maintenance. How it governs triad maintenance is not clear. We confirm interaction with BIN1, a key protein for T-tubule biogenesis, and also the increase in DNM2 levels that occurs with loss of MTM1. Interruption of the MTM1-BIN1 interaction, or overexpression of DNM2, may be sufficient to explain how MTM1 influences the triad, though the role of BIN1-MTM1 binding still requires further study, and the mechanism(s) by which DNM2 protein levels (but not RNA levels) are increased in XLMTM is unknown. An interesting alternative consideration is that DNM2/BIN1/MTM1 participate in a membrane recycling process that utilizes the endosome and that governs triad membrane turnover. Such as turnover process has yet to be elucidated in skeletal muscle, but it fits with the cellular functions of these proteins.

We again demonstrate that DNM2 protein levels are increased, show that they are increased at all time points tested, and reveal via RNA sequencing that DNM2 transcripts are not increased. This supports the importance of DNM2 overexpression in the disease process, and is consistent with the impressive impact of DNM2 lowering treatments on *Mtm1* KO mouse phenotypes.

In addition to DNM2 lowering, several other therapeutic strategies have been identified using the XLMTM mouse model. These include gene replacement therapy, enzyme replacement therapy, PIK3C2B lowering, and pathway modifiers such as rapamycin. Moving forward, it will be critical to have harmonization in the measures examined for existing in new treatments, so that the field can best compare and contrast these therapies. Lack of uniformity in terms of techniques and outcomes has been identified as an important problem in pre-clinical therapy development, and a key contributor to potential failures of translation for drugs in other rare diseases. We thus encourage use of standard set of experiments and techniques when evaluating treatments in the Mtm1 KO mouse model. From a general phenotype prospective, body weight, open field gait analysis, and survival are robust and reproducible. From a pathology prospective, mitochondrial organization, myofiber size, and triad number are easily quantified and reflect key aspects of the disease process. Molecular investigations should relate both to general disease pathway changes, such as DNM2 and polyubiquitin changes, and to the specific therapeutic, so as to demonstrate clear target engagement.

Lastly, our data highlight new potential avenues for investigation into XLTM disease pathomechanisms. This is highlighted, for example, when considering our longitudinal transcriptomics and subcellular proteomics, particularly in comparison to previous multi omic studies. For example, we identified increased expression of TUBB6 through both RNAseq and proteomics. This protein is of potential interest given that its persistent upregulation has been associated with disorganization of microtubule networks and return to a more immature developmental stage in skeletal muscle(Randazzo et al., 2019), both observations seen in XLMTM skeletal muscle.

## Conclusion

In summary, we present molecular and cellular changes of the XLMTM mouse model at four stages of the murine disease process (Figure 9). Our data should provide a roadmap for testing of future therapies and for elucidating disease pathomechanisms.

## Acknowledgements

This work was supported by research grants from CIHR, NSERC, and the Myotubular Trust, and by the Mogford Campbell Family Chair fund (JJD). We thank the Toronto Centre for Phenogenomics for assistance with mouse husbandry. We thank the Proteomics core facility at Mt Sinai (led by Anne Claude Gingras) for help with interpretation of IP-MS data. We acknowledge the Centre for Computational Medicine for assistance with analysis of RNA sequencing data, and The Centre for Applied Genomics (TCAG) for all sequencing that was performed.

## Methods

### Mouse husbandry and genotyping

Mice were housed at the Toronto Centre for Phenogenomics. The *Mtm1* KO line was originally generated at the IGBMC (Strasberg, France)(Buj-Bello et al., 2002). The line was rederived onto the C57BL6J background at the Toronto Centre for Phenogenomics. To limit genetic drift, carrier female mice were backcrossed to the original C57BL6J strain every 3-4 generations. All mice had unlimited access to water and food and were fed on regular rodent diet (Envigo 2918). Mice were euthanatized for tissue harvest by carbon dioxide chamber or cervical dislocation. All animal procedures were performed in compliance with the Animals for Research Act of Ontario and the Guidelines of the Canadian Council on Animal Care (AUP#22-0255H).

A PCR based method was used to detect exon 4 deletion and following primers were used:

Fwd: AATGTGTGCATGTTTGGACC

Rev: ACAGTGATGCACAGAGAGGA

### Histology and immunostaining

Frozen muscle tissue (tibialis anterior) was cut using a cryostat into 8-mm cross sections and then stained with Mayer’s hematoxylin and eosin (H&E) or succinate dehydrogenase (SDH). For indirect immunofluorescence (IF), frozen sections were blocked for 1 hour, then incubated overnight at 4C with Dysferlin antibody (catalog ab124684, Abcam), diluted 1:100 in blocking solution. Secondary antibodies (Alexa Fluor 488; Invitrogen) were applied for 1 hour at room temperature at 1:1,000 dilution. Micrographs were captured with an Infinity1 camera (Lumenera Corp.) through an Olympus BX43 light microscope. Quantification of the number of centrally nucleated fibers, myofiber size, SDH staining and Dysferlin IF was performed manually using Image J from H&E, SDH and Dysferlin IF photographs taken at 20x magnification.

### Transmission electron microscopy

Muscles (tibialis anterior) were fixed in 2% glutaraldehyde containing 0.1 M sodium cacodylate buffer overnight at 4 C, then taken to the Advanced Bioimaging Center (The Hospital for Sick Children, Toronto, Canada). Sections 90 nm thick were cut on an RMC MT6000 ultramicrotome, stained with uranyl acetate and lead citrate, and viewed with an FEI Tecnai 20 microscope.

### Western blot analysis

Muscles (TA, hamstring, quadriceps, gastrocnemius) were flash-frozen on dry ice and immediately stored at -80°C. Tissue was lysed with steel beads in TissueLyserII (Qiagen) and cells were lysed using cell scraper in the presence of RIPA buffer (supplemented with protease and phosphatase inhibitors) to extract protein. Samples were centrifuged at 12,700xg for 30 minutes at 4°C. Pierce™ BCA protein assay kit (Thermo Fisher) was used to quantify the protein levels in the supernatant. 30µg protein per lane was prepared with 4X loading buffer supplemented with DTT. Samples were boiled for 5 mins at 95°C with the exception of samples for RyR1 blots which were heated at 37°C for 10 mins. Samples were run on SDS-PAGE gels until necessary and then transferred on polyvinylidene fluoride (PVDF) membranes by semi-dry transfer method. Following the transfer, the membranes were treated with total-protein REVERT stain and visualized by LICOR instrument at 700nm. Membranes were blocked with 3% BSA for 1 hour at room temperature and then incubated with primary antibody overnight at 4°C. Primary antibodies used were DNM2 (1/1000, Santa Cruz sc-166669), PolyUbiquitinated proteins (1/1000, Sigma 04-262), p62 (1/1000, CST 5114S), acetylated α-tubulin (1/2000, Sigma T6793), α-tubulin (1/1000, CST 2144), MTM1 (Clone 2827, Dr. Jocelyn Laporte, IGBMC France). Secondary antibodies used were: horseradish peroxidase-conjugated goat anti-rabbit or goat anti-mouse IgG secondary antibody (1:5000, BioRad). Blots were visualized using chemiluminescence (Clarity Max™ ECL, BioRad or Western Lightning ECL Pro, PerkinElmer) at the Gel Doc™ XR with Gel Documentation System (BioRad). Membranes were stripped using Restore Western Blot Stripping Buffer (Thermo Scientific). Total protein and band intensities were measured using Fiji and relative expression of proteins were normalized to the total protein staining of the lane. Every blot included 4 biological replicates and was run in triplicates for technical replicates.

### RNA-seq and differential expression analysis

RNA extraction, library preparation, and RNA sequencing were performed as previously described(Maani et al., 2018). Briefly, RNA was extracted from WT (n=4) and *Mtm1* KO (n=4) tibialis anterior muscle at 14, 21 and 35 days by Qiagen Fibrous RNeasy kit in accordance with the manufacturer’s protocol. NEBNext® Ultra™ II Directional RNA Library Prep Kit for Illumina was used for library preparation and approximately 1000ng of RNA was used as starting material. The cut-off for RNA integrity number (RIN) was 8. Sequencing was done at The Centre for Applied Genomics (TCAG) using Illumina HiSeq 2500. Paired end read sequencing was performed to achieve 62-67 million reads per sample. Reads were aligned to the reference genome (GRCm38/MM10 version of the *Mus musculus* genome) and transcriptome using STAR51 two-step alignment to generate a Binary Alignment Map file (BAM file). Coordinate sorted BAM files were used to quantify transcript abundance (count data) using HTSeq52. Raw read counts generated were used as input for differential gene expression analysis, carried out using both DESeq2 and edgeR R/Bioconductor packages. This was carried out in a pairwise manner between any two conditions that were considered. FDR adjusted p-values from both edgeR and DESeq2 were used to determine genes that are significantly differentially expressed between the conditions tested. Metascape (Zhou et al., 2019) was used for GO term enrichment of differentially expressed genes (DEGs) between timepoints(Zhou et al., 2019). Finally, we compared our DEGs to those reported in similar mouse models previously(Djeddi et al., 2021).

### Subcellular proteomics

The methods used in sample preparation, namely choice of lysis buffers and centrifugation speeds, were based on previously published methods(Dimauro et al., 2012; Itzhak et al., 2016). First, 21 day TA muscle (n =3 per replicate) was cut into small sections over dry ice. A total of three biological replicates were used for each genotype. Next, sections were homogenized in a glass douncer over ice containing detergent-free lysis buffer (DFL) (250 mM sucrose, 50 mM HEPES buffer pH 7.4, 5mM MgCl2, 10 mM ATP, protease and phosphatase inhibitor cocktails). Tissue was homogenized with 10 strokes of loose pestle followed by 20 strokes of the tight pestle. Samples were then maintained on ice for 30 min.

Homogenates were decanted in fresh 2mL tubes and vortexed at max speed for 10s, and centrifuged at 200 x g for 5 min to remove large cellular debris and insoluble myofibrillar components. The pellet was discarded, and supernatant (S1) was moved into fresh 2mL tubes, vortexed at max speed for 10s, and centrifuged at 1000 x g for 15 min at 4°C. Supernatant (S2) was moved into fresh 2mL tubes while the “Nuclear” pellet (P1) was placed on dry ice. S2 was vortexed at max speed for 10s, and centrifuged at 800 x g for 10 min at 4°C to remove remaining/contaminating nuclei. Supernatant (S3) was moved into fresh 50 mL Nalgene round-bottom centrifuge tubes while the pellet (P2) containing residual nuclei was discarded. S3 was vortexed at max speed for 10s, and centrifuged at 24,000 x g for 20 min at 4°C. “Cytosolic” supernatant (S4) into new 2 mL conical tubes and placed on dry ice. Finally, the “Organellar/Membrane” pellet (P3) was placed on dry ice.

Lysis buffer (RIPA buffer + 1% SDS + protease and phosphatase inhibitors) was used to re-suspend pellets P1 and P3, heated to 72°C for 5 min with intermittent vortexing. Resuspended pellets and cytosolic fraction S3 were sonicated for 10 cycles (30s on/off) using a Diagenode Bioruptor at high setting and at 4°C. Nuclear samples (P1) were treated with 25U/mL benzonase endonuclease at RT for 30 min. Protein concentration of all samples were quantified using the BCA assay before being sent for proteomics analysis.

Additional sample preparation and mass spectrometry analysis were performed by the Network Biology Collaborative Centre (NBCC) at Lunenfeld-Tanenbaum Research Institute (Toronto, ON). 100ug of protein was processed from each sample using S-traps (ProtiFi) per manufacturer’s protocols and digested with 5 µg of trypsin each. Next, 10µg of each sample was labeled with 80µg of TMT 10-plex label per manufacturer protocols. For each subcellular fractionation (nucleus, membrane, cytoplasm), we used n=3 biological replicates from WT or *Mtm1* KO mutants. Each subcellular fraction was multiplexed with 6 TMT labels for three independent 6-plex experiments. Trapped and trypsin-digested peptides were acquired on a 180min gradient using an Orbitrap Fusion Lumos Tribrid mass spectrometer. Data was searched with SequestHT using the UP000000589 proteome (*Mus musculus*) generated by Uniprot, post-processed with Percolator, and analyzed using Proteome Discoverer 2.2.0.388 (Thermo Scientific).

Metascape was used for GO term enrichment and PPI network analysis(Zhou et al., 2019). To test whether or not the obtained fractions were enriched in expected proteins, we cross-referenced identified proteins to consensus lists from(Orre et al., 2019). This resource database predicts the subcellular location of proteins based on mass spectrometry data obtained from five cell lines and defines four enrichment neighborhoods: 1) cytosolic, 2) nuclear, 3) secretory, and 4) mitochondrial. Some proteins show consensus locations across cell lines where the protein was identified (e.g. All cell lines show protein X to be cytosolic). We restricted our enrichment analysis to refer only to proteins with consensus in 3 or more cell lines. Next, proteins identified in our fractions were cross-referenced to the consensus lists to determine their expected subcellular location. Peptide spectral matches (PSMs) were used as a proxy for relative abundance given that PSMs indicate how many times a peptide of the particular protein was selected for MS2 fragmentation and confidently identified. The percentage of PSMs for a given protein in each fraction was calculated over the total number of PSMs for that protein. In this way, we could visualize how often that peptide was identified in one fraction relative to the others to estimate the degree of our subcellular enrichment.

### Immunoprecipitation/Mass Spectrometry (IP-MS)

Muscle was harvested from 21 (n=3/genotype) and 36 day (n=2/genotype) old WT and *Mtm1*-6xHis mice and stored at -80C. Frozen muscle samples (»30mg) were minced and lysed in lysis buffer (50 mM Hepes-NaOH pH 8.0, 100 mM KCl, 0.1% NP40 and 10% glycerol, supplemented with 1 mM PMSF, 1 mM TCEP and 1X protease inhibitor cocktail (Sigma-Aldrich) using dounce tissue grinder (Wheaton Science, #357544). Samples were centrifuged at max speed for 30 minutes at 4°C and supernatants were collected. Magnetic dynabeads (ThermoFisher #10001D) were washed in PBS and incubated with 4μg of anti-6xHis antibody (Sigma-Aldrich #05-949) overnight at 4°C with gentle agitation. Supernatants were incubated with 25uL of pre-conjugated beads for 2 hours at 4°C with gentle agitation. Following this incubation, beads were washed once with lysis buffer, then washed twice times with wash buffer (20mM Tris-HCl (pH 8.0) and 2mM CaCl2). Beads were submitted to the NBCC, where samples were further washed, trypsin digested and eluted off the beads prior to liquid chromatography-mass spectrometry (LC-MS). SAINT analysis was performed by the facility.

### Data Statistical Analysis

All data presented are expressed as mean ± SEM (unless otherwise specified) and were analyzed for statistical significance by the unpaired two-tailed t-test (unless stated otherwise). P < 0.05 was considered to be statistically significant (*p<0.05, **p<0.01 and***p<0.001). GraphPad Prism software, version 8.0 for was used for generating graphs and performing statistical analysis.

## References

Al-Qusairi, L., Prokic, I., Amoasii, L., Kretz, C., Messaddeq, N., Mandel, J. L. and Laporte, J. (2013). Lack of myotubularin (MTM1) leads to muscle hypotrophy through unbalanced regulation of the autophagy and ubiquitin-proteasome pathways. FASEB J 27, 3384–3394.

Al-Qusairi, L., Weiss, N., Toussaint, A., Berbey, C., Messaddeq, N., Kretz, C., Sanoudou, D., Beggs, A. H., Allard, B., Mandel, J. L., et al. (2009). T-tubule disorganization and defective excitation-contraction coupling in muscle fibers lacking myotubularin lipid phosphatase. Proc Natl Acad Sci U S A 106, 18763–18768.

Amburgey, K., Tsuchiya, E., de Chastonay, S., Glueck, M., Alverez, R., Nguyen, C. T., Rutkowski, A., Hornyak, J., Beggs, A. H. and Dowling, J. J. (2017). A natural history study of X-linked myotubular myopathy. Neurology 89, 1355–1364.

Annoussamy, M., Lilien, C., Gidaro, T., Gargaun, E., Che, V., Schara, U., Gangfuss, A., D’Amico, A., Dowling, J. J., Darras, B. T., et al. (2019). X-linked myotubular myopathy: A prospective international natural history study. Neurology 92, e1852–e1867.

Bachmann, C., Jungbluth, H., Muntoni, F., Manzur, A. Y., Zorzato, F. and Treves, S. (2017). Cellular, biochemical and molecular changes in muscles from patients with X-linked myotubular myopathy due to MTM1 mutations. Hum Mol Genet 26, 320–332.

Beggs, A. H., Bohm, J., Snead, E., Kozlowski, M., Maurer, M., Minor, K., Childers, M. K., Taylor, S. M., Hitte, C., Mickelson, J. R., et al. (2010). MTM1 mutation associated with X-linked myotubular myopathy in Labrador Retrievers. Proc Natl Acad Sci U S A 107, 14697–14702.

Buj-Bello, A., Laugel, V., Messaddeq, N., Zahreddine, H., Laporte, J., Pellissier, J. F. and Mandel, J. L. (2002). The lipid phosphatase myotubularin is essential for skeletal muscle maintenance but not for myogenesis in mice. Proc Natl Acad Sci U S A 99, 15060–15065.

Carlier, R. Y. and Quijano-Roy, S. (2019). Myoimaging in Congenital Myopathies. Semin Pediatr Neurol 29, 30–43.

Childers, M. K., Joubert, R., Poulard, K., Moal, C., Grange, R. W., Doering, J. A., Lawlor, M. W., Rider, B. E., Jamet, T., Daniele, N., et al. (2014). Gene therapy prolongs survival and restores function in murine and canine models of myotubular myopathy. Sci Transl Med 6, 220ra210.

Cocanougher, B. T., Flynn, L., Yun, P., Jain, M., Waite, M., Vasavada, R., Wittenbach, J. D., de Chastonay, S., Chhibber, S., Innes, A. M., et al. (2019). Adult MTM1-related myopathy carriers: Classification based on deep phenotyping. Neurology 93, e1535–e1542.

Cowling, B. S., Chevremont, T., Prokic, I., Kretz, C., Ferry, A., Coirault, C., Koutsopoulos, O., Laugel, V., Romero, N. B. and Laporte, J. (2014). Reducing dynamin 2 expression rescues X-linked centronuclear myopathy. J Clin Invest 124, 1350–1363.

Cowling, B. S., Toussaint, A., Amoasii, L., Koebel, P., Ferry, A., Davignon, L., Nishino, I., Mandel, J. L. and Laporte, J. (2011). Increased expression of wild-type or a centronuclear myopathy mutant of dynamin 2 in skeletal muscle of adult mice leads to structural defects and muscle weakness. Am J Pathol 178, 2224–2235.

Dimauro, I., Pearson, T., Caporossi, D. and Jackson, M. J. (2012). A simple protocol for the subcellular fractionation of skeletal muscle cells and tissue. BMC Res Notes 5, 513.

Djeddi, S., Reiss, D., Menuet, A., Freismuth, S., de Carvalho Neves, J., Djerroud, S., Massana-Munoz, X., Sosson, A. S., Kretz, C., Raffelsberger, W., et al. (2021). Multiomics comparisons of different forms of centronuclear myopathies and the effects of several therapeutic strategies. Mol Ther 29, 2514–2534.

Dowling, J. J., Vreede, A. P., Low, S. E., Gibbs, E. M., Kuwada, J. Y., Bonnemann, C. G. and Feldman, E. L. (2009). Loss of myotubularin function results in T-tubule disorganization in zebrafish and human myotubular myopathy. PLoS Genet 5, e1000372.

Dowling, J. J., Weihl, C. C. and Spencer, M. J. (2021). Molecular and cellular basis of genetically inherited skeletal muscle disorders. Nat Rev Mol Cell Biol.

Fetalvero, K. M., Yu, Y., Goetschkes, M., Liang, G., Valdez, R. A., Gould, T., Triantafellow, E., Bergling, S., Loureiro, J., Eash, J., et al. (2013). Defective autophagy and mTORC1 signaling in myotubularin null mice. Mol Cell Biol 33, 98–110.

Gavriilidis, C., Laredj, L., Solinhac, R., Messaddeq, N., Viaud, J., Laporte, J., Sumara, I. and Hnia, K. (2018). The MTM1-UBQLN2-HSP complex mediates degradation of misfolded intermediate filaments in skeletal muscle. Nat Cell Biol 20, 198–210.

Gonorazky, H. D., Bonnemann, C. G. and Dowling, J. J. (2018). The genetics of congenital myopathies. Handb Clin Neurol 148, 549–564.

Gupta, V. A., Hnia, K., Smith, L. L., Gundry, S. R., McIntire, J. E., Shimazu, J., Bass, J. R., Talbot, E. A., Amoasii, L., Goldman, N. E., et al. (2013). Loss of catalytically inactive lipid phosphatase myotubularin-related protein 12 impairs myotubularin stability and promotes centronuclear myopathy in zebrafish. PLoS Genet 9, e1003583.

Hnia, K., Tronchere, H., Tomczak, K. K., Amoasii, L., Schultz, P., Beggs, A. H., Payrastre, B., Mandel, J. L. and Laporte, J. (2011). Myotubularin controls desmin intermediate filament architecture and mitochondrial dynamics in human and mouse skeletal muscle. J Clin Invest 121, 70–85.

Hnia, K., Vaccari, I., Bolino, A. and Laporte, J. (2012). Myotubularin phosphoinositide phosphatases: cellular functions and disease pathophysiology. Trends Mol Med 18, 317–327.

Itzhak, D. N., Tyanova, S., Cox, J. and Borner, G. H. (2016). Global, quantitative and dynamic mapping of protein subcellular localization. Elife 5.

Jungbluth, H., Treves, S., Zorzato, F., Sarkozy, A., Ochala, J., Sewry, C., Phadke, R., Gautel, M. and Muntoni, F. (2018). Congenital myopathies: disorders of excitation-contraction coupling and muscle contraction. Nat Rev Neurol 14, 151–167.

Ketel, K., Krauss, M., Nicot, A. S., Puchkov, D., Wieffer, M., Muller, R., Subramanian, D., Schultz, C., Laporte, J. and Haucke, V. (2016). A phosphoinositide conversion mechanism for exit from endosomes. Nature 529, 408–412.

Laporte, J., Hu, L. J., Kretz, C., Mandel, J. L., Kioschis, P., Coy, J. F., Klauck, S. M., Poustka, A. and Dahl, N. (1996). A gene mutated in X-linked myotubular myopathy defines a new putative tyrosine phosphatase family conserved in yeast. Nat Genet 13, 175–182.

Lawlor, M. W. and Dowling, J. J. (2021). X-linked myotubular myopathy. Neuromuscular Disorders.

Liu, N., Bezprozvannaya, S., Shelton, J. M., Frisard, M. I., Hulver, M. W., McMillan, R. P., Wu, Y., Voelker, K. A., Grange, R. W., Richardson, J. A., et al. (2011). Mice lacking microRNA 133a develop dynamin 2-dependent centronuclear myopathy. J Clin Invest 121, 3258–3268.

Maani, N., Sabha, N., Rezai, K., Ramani, A., Groom, L., Eltayeb, N., Mavandadnejad, F., Pang, A., Russo, G., Brudno, M., et al. (2018). Tamoxifen therapy in a murine model of myotubular myopathy. Nat Commun 9, 4849.

McEntagart, M., Parsons, G., Buj-Bello, A., Biancalana, V., Fenton, I., Little, M., Krawczak, M., Thomas, N., Herman, G., Clarke, A., et al. (2002). Genotype-phenotype correlations in X-linked myotubular myopathy. Neuromuscul Disord 12, 939–946.

Randazzo, D., Khalique, U., Belanto, J. J., Kenea, A., Talsness, D. M., Olthoff, J. T., Tran, M. D., Zaal, K. J., Pak, K., Pinal-Fernandez, I., et al. (2019). Persistent upregulation of the beta-tubulin tubb6, linked to muscle regeneration, is a source of microtubule disorganization in dystrophic muscle. Hum Mol Genet 28, 1117–1135.

Reumers, S. F. I., Braun, F., Spillane, J. E., Bohm, J., Pennings, M., Schouten, M., van der Kooi, A. J., Foley, A. R., Bonnemann, C. G., Kamsteeg, E. J., et al. (2021). Spectrum of Clinical Features in X-Linked Myotubular Myopathy Carriers: An International Questionnaire Study. Neurology 97, e501–e512.

Royer, B., Hnia, K., Gavriilidis, C., Tronchere, H., Tosch, V. and Laporte, J. (2013). The myotubularin-amphiphysin 2 complex in membrane tubulation and centronuclear myopathies. EMBO Rep 14, 907–915.

Sabha, N., Volpatti, J. R., Gonorazky, H., Reifler, A., Davidson, A. E., Li, X., Eltayeb, N. M., Dall’Armi, C., Di Paolo, G., Brooks, S. V., et al. (2016). PIK3C2B inhibition improves function and prolongs survival in myotubular myopathy animal models. J Clin Invest 126, 3613–3625.

Tasfaout, H., Buono, S., Guo, S., Kretz, C., Messaddeq, N., Booten, S., Greenlee, S., Monia, B. P., Cowling, B. S. and Laporte, J. (2017). Antisense oligonucleotide-mediated Dnm2 knockdown prevents and reverts myotubular myopathy in mice. Nat Commun 8, 15661.

Volpatti, J. R., Al-Maawali, A., Smith, L., Al-Hashim, A., Brill, J. A. and Dowling, J. J. (2019). The expanding spectrum of neurological disorders of phosphoinositide metabolism. Dis Model Mech 12.

Zhou, Y., Zhou, B., Pache, L., Chang, M., Khodabakhshi, A. H., Tanaseichuk, O., Benner, C. and Chanda, S. K. (2019). Metascape provides a biologist-oriented resource for the analysis of systems-level datasets. Nat Commun 10, 1523.

